# Antibiotic treatment modestly reduces protection against Mycobacterium tuberculosis reinfection in macaques

**DOI:** 10.1101/2023.12.19.570845

**Authors:** Sharie Keanne Ganchua, Pauline Maiello, Michael Chao, Forrest Hopkins, Douaa Mugahid, Philana Ling Lin, Sarah M. Fortune, JoAnne L. Flynn

## Abstract

Concomitant immunity is generally defined as an ongoing infection providing protection against reinfection^1^. Its role in prevention of tuberculosis (TB) caused by *Mycobacterium tuberculosis* (Mtb) is supported by epidemiological evidence in humans as well as experimental evidence in mice and non-human primates (NHPs). Whether the presence of live Mtb, rather than simply persistent antigen, is necessary for concomitant immunity in TB is still unclear. Here, we investigated whether live Mtb plays a measurable role in control of secondary Mtb infection. Using cynomolgus macaques, molecularly barcoded Mtb libraries, PET-CT imaging, flow cytometry and cytokine profiling we evaluated the effect of antibiotic treatment after primary infection on immunological response and bacterial establishment, dissemination, and burden post-secondary infection. Our data provide evidence that, in this experimental model, treatment with antibiotics after primary infection reduced inflammation in the lung but was not associated with a significant change in bacterial establishment, dissemination or burden in the lung or lymph nodes. Nonetheless, treatment of the prior infection with antibiotics did result in a modest reduction in protection against reinfection: none of the 7 antibiotic treated animals demonstrated sterilizing immunity against reinfection while 4 of the 7 non-treated macaques were completely protected against reinfection. These findings support that antibiotic-treated animals were still able to restrict bacterial establishment and dissemination after rechallenge compared to naïve macaques, but not to the full extent of non-antibiotic treated macaques.

## Introduction

Concomitant immunity is the ability of an ongoing infection to protect against subsequent infection and has been considered a hallmark of tuberculosis^2^ and leishmaniasis^3^. The most striking recent evidence for that in TB was reported in a meta-analysis of TB cases in healthcare workers by Andrews and colleagues^2^. In that study, the authors used data from approximately 20,000 individuals from 23 prospective cohorts to estimate whether there was a measurable difference in the incidence rate of symptomatic TB disease in healthcare workers with latent TB infection (LTBI; defined as PPD positive at baseline) and those without (PPD negative at baseline). They found that LTBI subjects had a 79% lower risk of developing active TB compared to an uninfected person upon secondary exposure. In other words, prior infection with Mtb seems to significantly reduce the chance of developing active TB disease. While the conclusion is intriguing, in these studies the timing of primary infection was unknown, and it is unclear whether the LTBI individuals were carriers of live Mtb, had cleared the initial infection, or if protection was due to long-term immunological reprogramming that correlates with PPD positivity^4^. There was also no definitive evidence that the patients were re-exposed to or infected with Mtb during the course of each study, so we cannot exclude the possibility that TB disease could have been due to reactivation of a latent infection, at least in some cases. Finally, it is not possible to determine whether subjects were reinfected and contained the secondary infection, since the only outcome measure in these studies was development of active TB.

Indeed, one recent study of recurrent TB in a high TB burden region in Southern China showed that 75% of cases were due to reactivation compared to 25% due to exogenous reinfection^5^. If generalizable across populations, these findings suggest that reinfection does occur in humans but is the minority of cases, complicating the interpretation of epidemiological data in the context of understanding concomitant immunity. Other studies from areas with high levels of endemic TB^6,7^ reported the detection of multiple Mtb strains in the sputum of both HIV negative and positive individuals providing stronger and direct evidence that reinfection occurs in humans. The small numbers make it difficult to conclude with certainty whether HIV increases the rate of reinfection, but they do report a higher proportion of HIV positive patients carrying multiple Mtb strains, potentially implicating CD4+ T cells or other immune factors as important players in restricting reinfection with Mtb. In general, the immunological changes associated with protection against reinfection remains largely unexplored.

Another open question in the field is whether live Mtb bacilli are necessary for concomitant immunity. There is evidence for that in mice as demonstrated by a study where mice were treated with antibiotics before reinfection with Mtb. These animals had 10-1,000-fold fewer Mtb bacilli in their lungs 30 days post-infection, compared to untreated controls. However, protection waned and lung bacterial burden increased to the level of control animals by 60 days post-infection^8^. More recently, a study in mice demonstrated that a contained intradermal Mtb infection in the ear and ear-draining lymph node resulted in reduced lung bacterial burden following rechallenge by aerosol^9^. This protective effect was significantly reduced when intradermally infected animals were treated with antibiotics prior to rechallenge, suggesting that a live ongoing infection is important for concomitant immunity in small mammals.

It has been difficult to determine whether the elimination of live bacteria after primary infection substantially impairs concomitant immunity in humans. According to studies in China^10^ and Spain^11^ the incidence rate of recurrent TB disease after successful treatment is 18 and 14.6 times the incidence rate of initial TB disease in the general population, respectively. A study in South Africa which used DNA fingerprinting methods to discriminate between reactivation and reinfection showed a fourfold higher incidence rate of active TB disease after successful antibiotic treatment compared to the general population^12^. Contrary to the findings in mice, and despite the large range in disease incidence rates, these numbers suggest that treatment from prior infection is associated with a higher likelihood of reinfection, or at least development of active TB after re-exposure, in humans. A potential explanation for this discrepancy between human and mouse studies include selection bias towards individuals with predisposition to Mtb infection and development of symptomatic TB disease in the clinical research setting, that symptomatic TB disease in humans increases the susceptibility of an individual to subsequent Mtb infection and disease upon subsequent exposure, and/or that elimination of live Mtb resulting from drug treatment increased susceptibility of these individuals to reinfection and active TB disease.

To address the limitations faced in clinical research settings, we previously investigated concomitant immunity in a non-human primate model, namely cynomolgus macaques^13^. In that study, primary and secondary infections were administered to macaques in a controlled setting and each infection was distinguishable using genomic barcodes unique to each strain. The results demonstrated the formation of fewer granulomas and a 10,000-fold reduction in bacterial burden from the second infection in macaques with ongoing primary infection, providing strong evidence for the occurrence of concomitant immunity in primates upon Mtb reinfection. Furthermore, protection was not associated with the degree of primary disease. Given the robust protection against reinfection, it was not possible to identify correlates or mechanisms of protection associated with concomitant immunity.

In this study, we investigated the relative importance of live bacteria for concomitant immunity in macaques. Our results show that antibiotic treatment after primary infection was effective in greatly reducing Mtb bacterial burden but only had a modest impact on protection against secondary infection as reflected by bacterial establishment, dissemination, or burden. We also observed no difference in T cell numbers or functions by flow cytometry after antibiotic treatment and prior to reinfection, although there were changes in the concentration of some cytokines in the airways, several of which are typically thought to be produced by innate immune cells. Overall, this study demonstrates the utility of this model for studying concomitant immunity in NHPs since antibiotic treatment of prior infection does not abolish protection against reinfection..

### Study design

We previously demonstrated that concurrent infection with Mtb prevented the establishment of a secondary infection in cynomolgus macaques (*Macaca fascicularis*)^13^. In this study, we sought to determine whether the persistence of live Mtb after the primary infection was required for protection against reinfection.

Fourteen cynomolgus macaques were first infected with a barcoded library of Mtb strain Erdman (Library P, ∼15 CFU; **Figure 1A**). Serial ^18^F-fluorodeoxyglucose (FDG) PET-CT scans were performed monthly to monitor the progress of infection. Specifically, we tracked inflammation in both lung and lymph nodes, as well as the formation of granulomas or other lung pathologies. Standard anti-TB drug treatment (HRZE) was initiated in 7 animals (Abx group) 16 weeks post infection with library P and continued for 12 weeks to eliminate or greatly reduce bacterial load (represented as dark green in **Figure 1** and subsequent figures). The other 7 animals were not treated with HRZE (No-Abx group). The Abx animals were then rested for 1 month after completion of their antibiotic regimen, after which animals were re-infected with ∼10 CFU of Mtb (Erdman strain; Library S). Since the genome of each Mtb bacillus in Library P and S are uniquely barcoded, we can identify each bacterium’s library of origin, as well as trace bacterial dissemination from the lung to other tissues. Given that previous experiments supported that most granulomas are established by a single bacterium^14^, and the probability of our *in vivo* inoculum (∼10 CFU) containing the same barcoded bacteria more than once is <2%, we assume that the presence of the same barcode in two different sites is the result of dissemination as opposed to the presence of the same barcode in the original pool.

**Figure 1.**
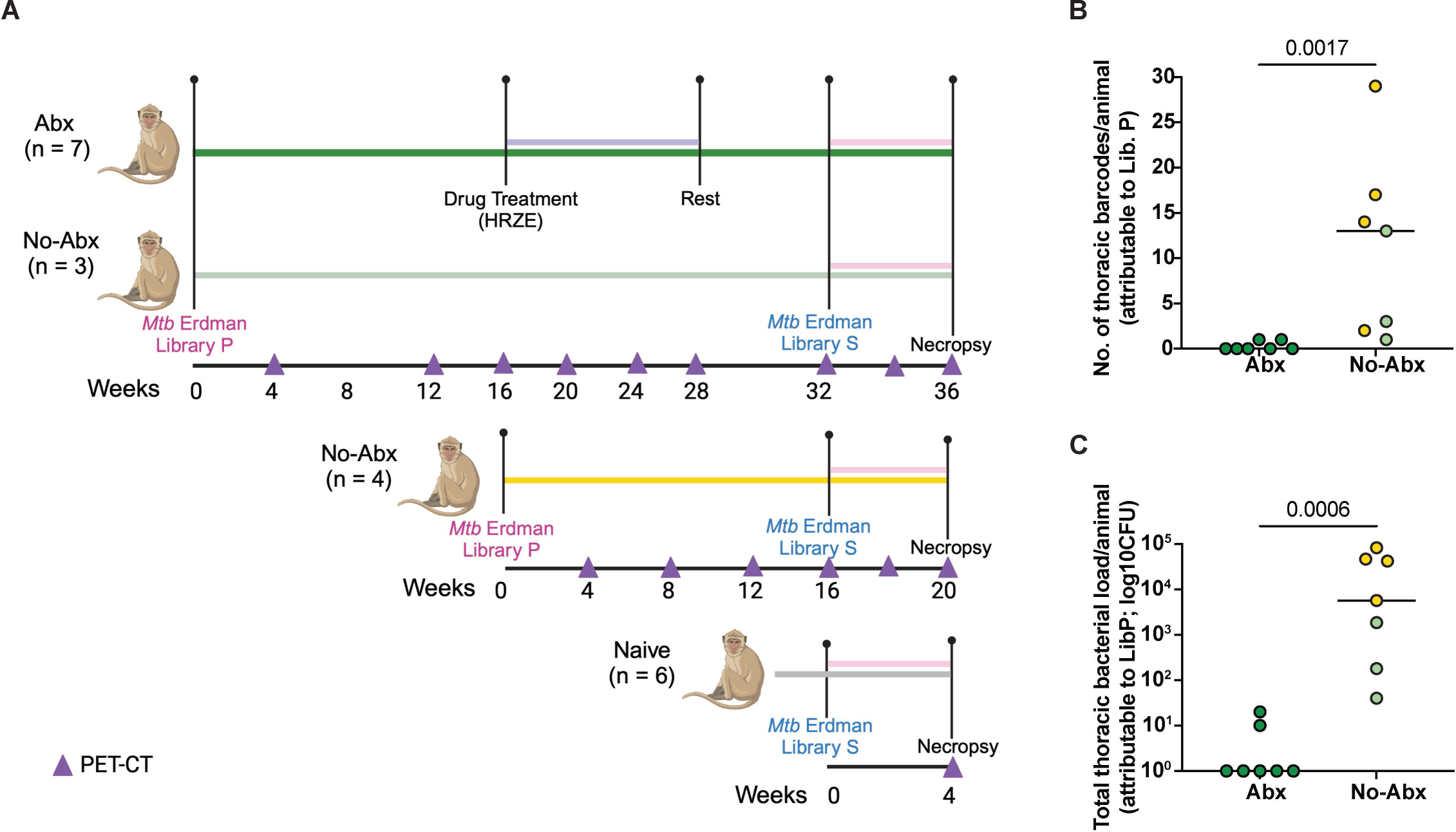
Experimental design and efficacy of antibiotic treatment. (**A**) Ten cynomolgus macaques were infected with Mtb Erdman Library P and followed for 32 weeks prior to re-challenge with Library S (Shown with dark green and light green solid lines). Seven animals were treated with antibiotics (Abx, purple line) 16 weeks after primary infection with Mtb Erdman Library P while three that followed the same timeline received no Abx before secondary infection. An additional cohort of four No-Abx animals were infected for 16 weeks prior to secondary infection (No-Abx, yellow). Both No-Abx groups were combined in subsequent analyses. A cohort of six naïve control animals were infected once with Library S four weeks before they were sacrificed; i.e. no primary infection (grey). The pink line corresponds to the period of secondary infection before necropsy. Triangles indicate time at which PET CT scans were performed. (**B**) Abx animals had fewer thoracic bacterial barcodes attributable to primary infection at necropsy compared to No-Abx animals, with a median count of 0 and 13, respectively. (**C**) Total thoracic bacterial burden attributable to primary infection (Library P) was greatly reduced in Abx animals compared to no-Abx animals with a median CFU of 1 and 5670, respectively. P-values are calculated based on a Mann-Whitney test, n=7 for both groups.

In the case of No-Abx animals, 3 were maintained on the same timeline as the drug treated animals, i.e. secondary infection occurred 32 weeks after primary infection (represented as light green in **Figure 1** and subsequent figures). For this group, we chose macaques with relatively low levels of disease after primary infection based on PET CT scans, since they might otherwise not have survived long enough for reinfection. Four additional untreated macaques were rechallenged at 16 weeks post-Library P infection, recapitulating our previously published studies^13^ (represented in yellow in **Figure 1** and subsequent figures). A group of 6 animals naïve to prior infection were also infected with Library S. All animals in this study were necropsied 4-5 weeks post-secondary infection with Library S. Using the final PET CT scan as a map, we identified all granulomas present before and after secondary infection and harvested granuloma and other lung pathologies, all thoracic and peripheral lymph nodes, and uninvolved lung tissue. These tissues were processed individually for microbiological, cellular, and molecular analysis as described in the methods.

### Antibiotic treatment effectively reduces Mtb burden after primary infection

To determine whether antibiotic treatment was effective in reducing bacterial burden post-primary infection, we analyzed changes in total lung FDG activity during drug treatment as measured by serial PET CT scans. There was variability in reduction of lung FDG activity among the Abx animals, but on average there was a 70% reduction in lung inflammation comparing pre-and post-drug time points (**Figure S1A**). Though not observed in all animals in the Abx group, we took this as an indication that animals were responding to Mtb treatment. In contrast there was minimal change in lung FDG activity in the untreated animals during that same period (**Figure S1A**). We also sequenced the genomic barcodes of the Mtb strains that grew on 7H11 plates from each tissue sample at necropsy, using the PET CT scans as a map for obtaining each individual lesion and all lymph nodes and lung tissues. The Mtb library of origin was distinguishable by virtue of the different Library IDs integrated into the genomes of Libraries P and S, respectively. CFU and barcodes tracible to the primary infection were completely absent in five of the seven Abx animals with a median thoracic CFU of 15 and a median of 0 positive tissues per animal (**Figure 1B, C**). In comparison, all seven animals in the no-Abx group were positive for Library P barcodes, with a median thoracic CFU of 5670 and 13 thoracic barcodes per animal (**Figure 1B, C**; **Supplementary table 1 and 2**). It is worth noting that bacterial burden was higher in the animals infected with Library P for 16 weeks (yellow) versus 32 weeks (light green), consistent with previous observations of higher CFU per granuloma at earlier time points post-infection^14^. We concluded that antibiotic treatment was effective in substantially or completely reducing the bacterial load in macaques, since 5 out of 7 animals were completely sterile and the other two had lower CFU than any of the untreated animals.

### Antibiotic treatment has a modest effect on bacterial establishment, restriction, or dissemination upon secondary infection

We next sought to determine whether antibiotic treatment, and thereby a substantial reduction in live Mtb post-primary infection, affected Mtb establishment and bacterial burden after secondary infection; in other words whether live bacteria were necessary for concomitant immunity. For that, we quantified the number of barcodes at necropsy from the tissues that contained Library S, used for secondary infection. There was a significant reduction in the total number of unique barcodes from Library S in both the Abx and no-Abx groups compared to naïve animals with a median count of ∼ 1, 3, and 12, respectively (p<0.05, Dunn’s multiple comparisons, **Figure 2A**). However, there was no significant difference between the Abx and no-Abx groups. These results suggest that antibiotic treatment of a primary infection is still associated with a restriction in Mtb establishment in the host upon secondary infection.

**Figure 2.**
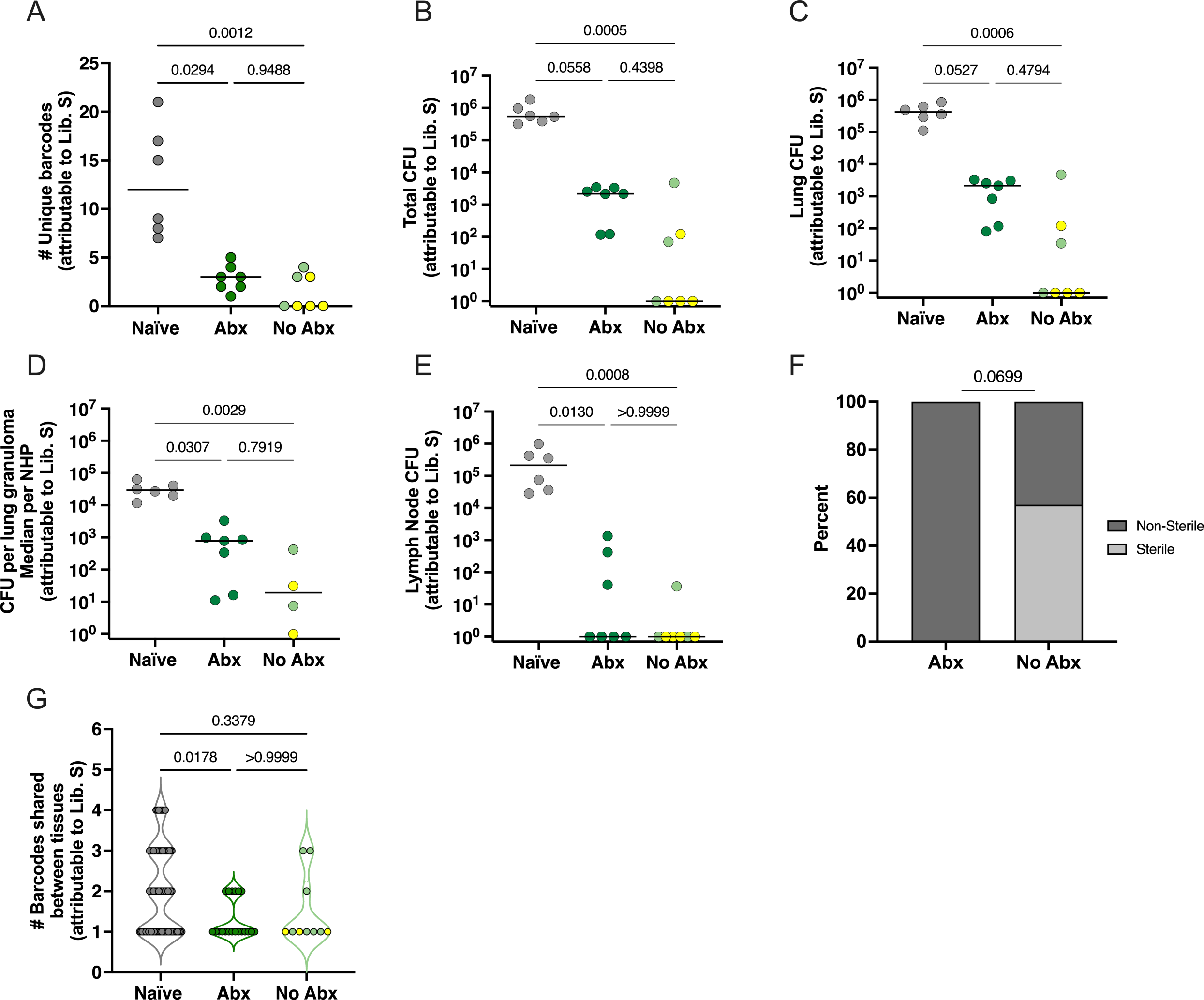
Antibiotic treatment prior to reinfection does not significantly affect bacterial establishment, load, or dissemination after secondary infection. (**A**) Number of unique bacterial barcodes from Library S quantified in tissues from reinfected or naïve animals. (**B-D**) Total thoracic CFU, lung CFU, and median CFU per granuloma per animal (note: 3 of the No Abx animals did not have granulomas so are not represented in this graph) (**E**) Thoracic lymph node CFU in naïve, Abx treated and no-Abx animals. (**F**) Distribution of sterile (no Library S CFU) animals in Abx and no-Abx treatment groups. (**G**) The number of barcodes shared between the lung and other tissues reflects bacterial dissemination from the lung; only the 3 No Abx animals with Library S barcodes are represented in this graph. Naïve, n=6; Abx, n=7; no Abx, n=7. (**A-E**) Each symbol is an animal; lines represent medians. Each symbol in **G** is a barcode. p-values are calculated based on Dunn’s multiple comparison testing post a Kruskal-Wallis test (**A-E**, **G**) and Fisher’s exact test (**F**).

We then quantified bacterial burden attributable to Library S in thoracic tissues including lung tissue, granulomas, and lymph nodes from all animals. Total thoracic bacterial burden was significantly lower in no-Abx animals compared to naïve controls (p=0.0005, Dunn’s multiple comparisons test) with a trend toward lower bacterial burden in Abx macaques compared to naïve controls (p=0.0558, Dunn’s test; **Figure 2B**). There was no significant difference comparing Abx and no-Abx macaques (p=0.4398). Similar results were observed when assessing Library S CFU in total lung, lung granuloma, or thoracic lymph nodes separately (**Figure 2C-E**). Based on these data, we concluded that antibiotic treatment of primary infection did not significantly reduce protection against secondary challenge. That said, 4 of the 7 no-Abx macaques had no detectable thoracic CFU attributable to Library S (i.e. 57% were sterile for Library S), while Library S CFU were recovered from all the Abx macaques (0% sterile) (p=0.0699 Fisher’s exact test, **Figure 2F**). These findings support a modest impact of antibiotic treatment post primary infection on reducing bacterial load after reinfection.

To quantify whether bacterial dissemination from the lung to other tissues was affected by antibiotic treatment, we compared the percentage of barcodes shared between the lung and other tissues. We observed no statistically significant difference between the Abx and no-Abx group (P>0.999; Dunn’s multiple comparisons test; **Figure 2G**).

### Antibiotic treatment reduces proinflammatory chemokines in airways but has no measurable effect on lymphocytes

We investigated whether antibiotic treatment resulted in measurable changes in immune responses prior to secondary infection. The lung FDG activity by PET indicated a 70% reduction on average in lung inflammation during drug treatment (**Figure S1**). To determine whether there was a difference in the airway immune environment pre-second infection in both groups, we examined lymphocyte population numbers, proliferation and cytokine responses in bronchoalveolar lavage (BAL) samples stimulated with ESAT-6 and CFP-10 obtained prior to secondary challenge. We found no difference in the total number of CD4+ or CD8+ T-cells (**Figure 3A-B**) comparing the Abx and no-Abx groups at 32 weeks post-primary Mtb infection. Furthermore, when isolated BAL cells were stimulated with ESAT-6 and CFP-10, there were no differences in the frequency of either CD4+ and CD8+ T-cells expressing IFN-γ, IL2, IL10, TNF, or Ki67 between Abx and no-Abx groups (**Figure 3C-L**).

**Figure 3.**
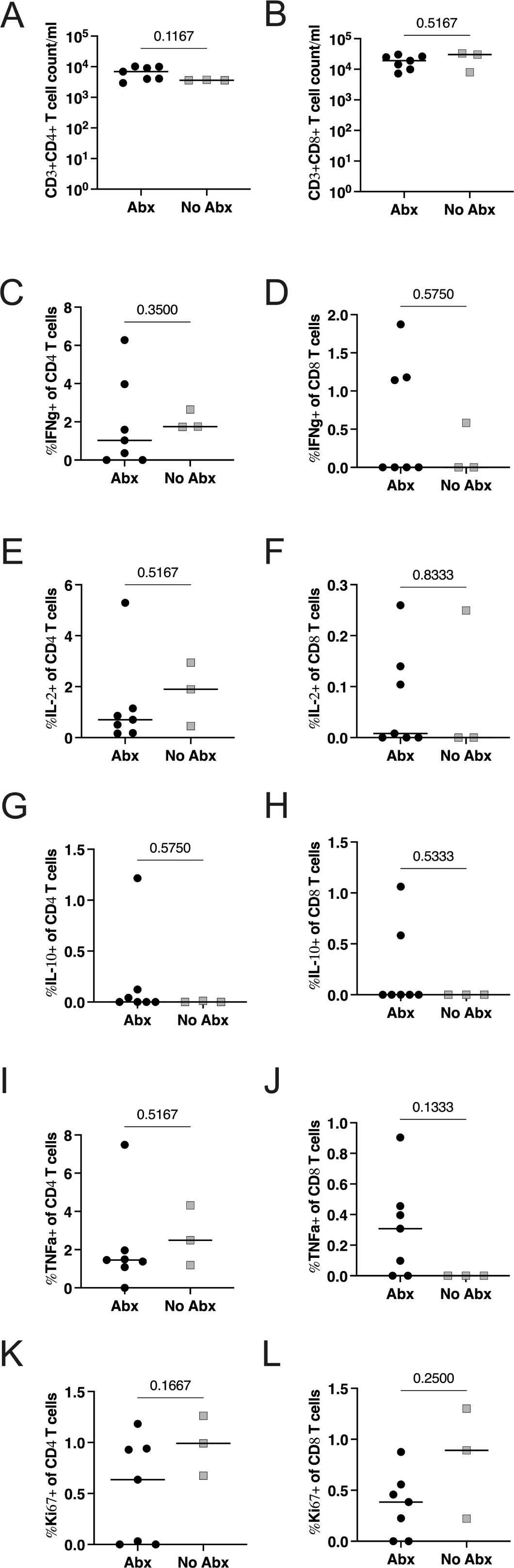
The effect of antibiotic treatment on CD3+ lymphocytes in the BAL prior to reinfection. (**A, B**) Number of CD4+ or CD8+ T-cells in the BAL. Frequency of BAL CD4+ and CD8+ T-cells producing pro-inflammatory cytokines IFNγ (C,D), IL2 (E, F), IL-10 (G, H), or TNF (I, J). Frequency of proliferating (Ki67) T cells (K, L). Median values depicted; p-values based on a Mann-Whitney test. Abx, n=7; no-Abx, n=3 (from the 32-week post-primary infection group).

We assessed BAL fluid for cytokines and chemokines using Luminex assays where we found significantly higher concentrations of pro-inflammatory chemokines, MIG (CXCL9), BLC (CXCL13), and MIP-1β (CCL4) in the no-Abx group compared to the Abx group (**Figure 4A-C)**. The levels of IL-1β and I-TAC were not significantly different (**Figure 4D-E**).

**Figure 4.**
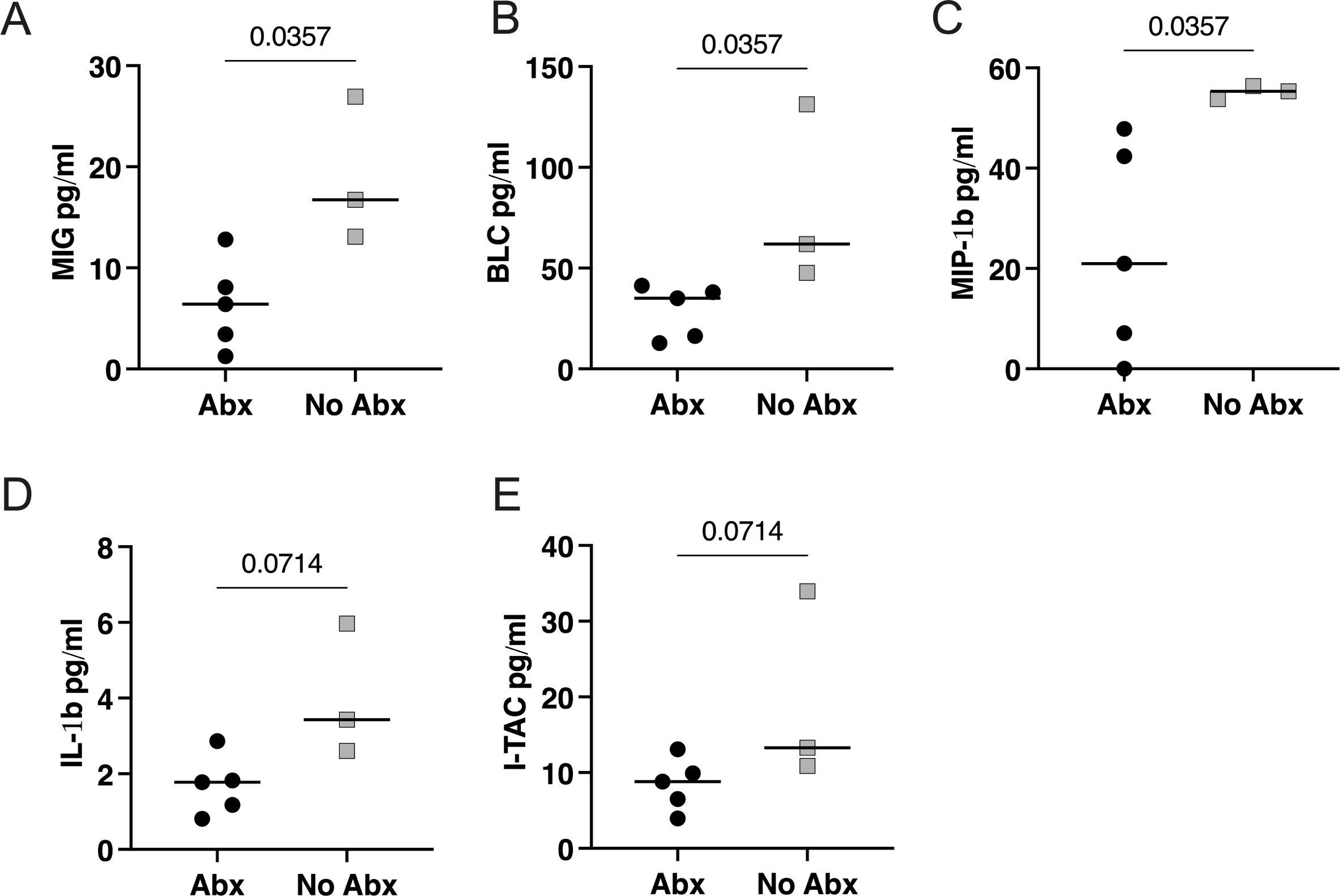
The effect of antibiotic treatment on cytokine and chemokine production in BAL fluid prior to reinfection. (**A**-**E**) Chemokines or cytokines in BAL fluid prior to reinfection as measured by Luminex. Median values depicted; p-values based on a Mann-Whitney test. Abx, n=7; no-Abx, n=3 (from the 32-week post-primary infection group).

Finally, there was no difference in the frequency of CD4+ or CD8+ T-cells (**Figure 5A-B**) in Library S positive granulomas from Abx or no-Abx macaques. The frequency of T cells producing CD107a, IFNγ+TNF, IL-2, IL-10, or IL-17 in granulomas was also not significantly different (**Figure 5C-L**). These results suggest that the immune environment in the airway (pre-second infection) and the granulomas from the second infection was similar between the Abx and no-Abx groups in terms of the measured immune variables, except for a change in a subset of pro-inflammatory chemokines in the airway.

**Figure 5.**
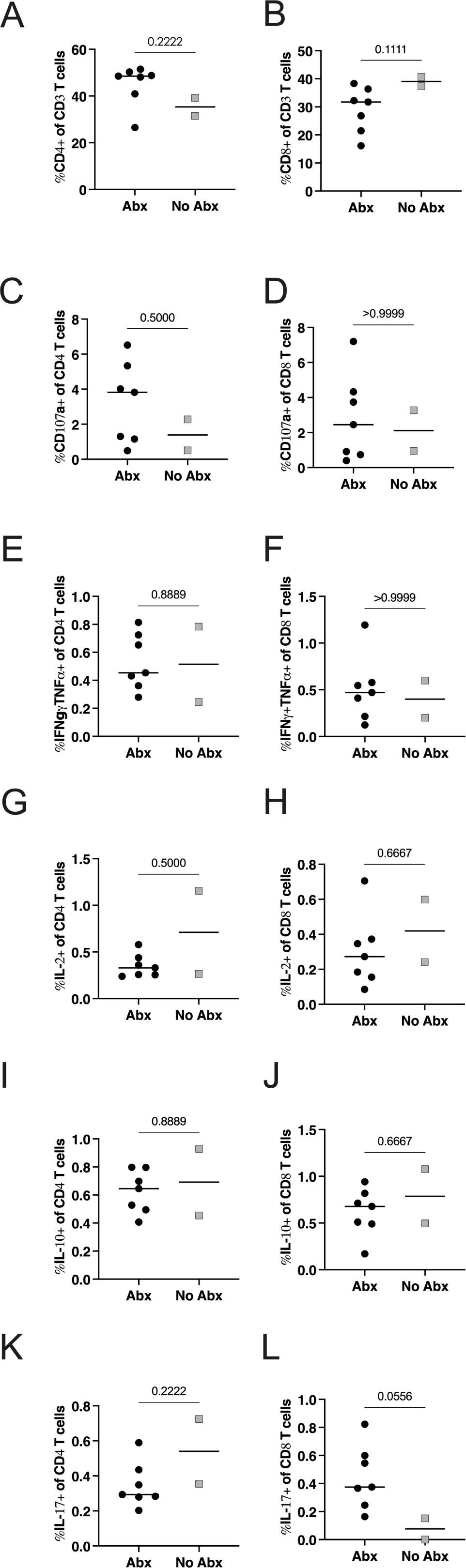
The effect of antibiotic treatment on CD3+ lymphocytes in granulomas arising post reinfection (attributable to Library S). (**A-B**) Frequency of CD4+ or CD8+ T cells in granulomas attributable to Library S from antibiotic treated versus untreated animals. (**C-L**) Frequency of CD4+ or CD8+ cells expressing effector molecules in library S granulomas. Median values depicted; p-values based on a Mann-Whitney test. Abx, n=7; no-Abx, n=2 (from the 32-week post-primary infection group).

## Discussion

Concomitant immunity against Mtb has been reported in humans based on epidemiological studies in humans with LTBI who have a presumed secondary exposure to Mtb^2^ and concomitant immunity protects against reinfection in mice and non-human primates (NHPs)^9,13^ and has been investigated in computational models^15^. However, epidemiologic data also indicate that humans previously treated for active TB have an elevated risk of developing a second case of TB^10,11^. Here we investigated whether live Mtb from an initial infection is necessary for protection against secondary infection in a NHP model. Our data indicate that treatment of a primary infection with antibiotics still provides protection against reinfection. All data considered, our findings suggest a modest reduction in protection due to antibiotic treatment, even if not statistically significant based on the number of animals used in this study.

Specifically, fewer Mtb bacilli successfully established infection in both antibiotic treated and untreated macaques compared to naïve animals and total thoracic bacterial burden, bacterial burden in the lung and in granulomas was similar between antibiotic treated and untreated animals. However, in the untreated group, there were more animals in which no bacteria from the second infection were found and median total CFU was lower compared to the antibiotic treated groups, though the difference was not statistically significant. Together, these data suggest a modest effect of antibiotic treatment on protection against reinfection. Although T cell numbers and functions in airways and granulomas were similar between the two reinfection groups, pro-inflammatory chemokines BLC (CXCL13), MIP-1β (CCL4), and MIG (CXCL9), which can participate in recruiting lymphocytes, were higher in the airways of untreated animals^16–18^.

Concomitant immunity is generally thought to rely on an ongoing or persistent infection. Although antibiotic treatment in this study was clearly effective at eliminating most, and in the majority of animals all, of the culturable Mtb, we speculate that persisting antigen may be responsible for the protection achieved with antibiotic treatment. We showed previously that macaques vaccinated with BCG delivered intravenously had robust protection against Mtb 6 months after vaccination even though the BCG was eliminated within a couple months of vaccination^19^. That robust protection may also rely in part on persistent mycobacterial antigens from BCG. An alternative hypothesis is that trained innate immunity, such as T cell “training” of macrophages, plays a role in immune protection in the absence of Mtb antigen as suggested by the increase in some chemokines produced by these cells. We have also recently reported evidence of altered myeloid cell state post-IV-BCG^20^. Finally, it is possible that the mechanisms of protection in antibiotic-treated and non-antibiotic treated animals are different. Ongoing effector T cells in lungs may be most relevant in the animals with concurrent Mtb infection, while in the antibiotic treated group, memory T cells may be playing a bigger role. We cannot distinguish among these possibilities in the current study.

Another factor to consider is the timing of the second infection. In C57BL/6 mice, protection from the second infection after drug treatment only lasted for 30 days. On day 60, the protection waned resulting in the bacterial burden of reinfected mice matching those of the controls (one infection only)^8^. Human epidemiology studies also support the theory of waning protection after successful treatment. A study in Uganda showed that in both HIV-negative and HIV-positive individuals, TB recurrence due to relapse was more common and occurred early at 6.5 months after completion of drug treatment while TB recurrence due to reinfection occurred much later at 14.2 months post-drug treatment^21^. Similarly, in Cape Town, South Africa, reinfection accounted for only 20% (9/44) of recurrent cases in the first year after drug treatment completion. This increased to 66% (57/86) after the first year. This study found a strong association between relapse and earlier occurrence ^22^. However, one study that analyzed extensive data from clinics in Cape Town, South Africa, found that the rate of Mtb reinfection and disease was highest in the first months after successful treatment. This rate rapidly declined by 2-3 years post-drug treatment and continued to decline until it reached the incidence rate of the general population at 10 years post-drug treatment^12^. Whether the protection we observed in Abx-treated animals will wane over time should be addressed in the future.

One limitation to this study was our bias in selecting the animals that received Abx treatment post primary infection. This was necessary since approximately half of Mtb infected cynomolgus macaques will develop active progressive TB disease. Consequently, we selected the 3 animals in the 32 week infection group that showed stable and low-level Library P infections as indicated by their serial PET CT scans to not receive Abx treatment. Furthermore, our no-Abx group is a combination of animals infected with Library P for 16 and 32 weeks prior to secondary infection which likely caused some heterogeneity post-secondary infection especially since we observed different CFU levels post primary infection (**Figure 1C**). This is in line with reducing animal numbers according to the 3Rs of animal study design (replacement, reduction, and refinement). That said, when we combined the two no-Abx groups for analysis, we noted no statistically significant differences between the animals using our primary outcome measure of total thoracic CFU attributable to Library S, albeit with low sample sizes in that comparison (Mann-Whitney test). Based on the Library S Total CFU (log_10_) variability in the non-drug treated group in this study, we performed a power analysis to determine how many animals would be needed to detect a statistically significant difference between drug treated and non-drug treated animals. This analysis indicated that we would need 17 animals per group. So while we suspect that drug treatment reduces protection against reinfection, our study was not powered to detect a meaningful difference between drug-treated and non-drug-treated animals. In addition, our immunologic measures were performed only in the macaques infected with Library P for 32 weeks and thus our power for statistical analyses was limited.

In summary, this study demonstrates that antibiotic treatment after primary infection does not significantly increase susceptibility to Mtb infection and disease after reinfection in macaques, but likely protection is not as robust as in non-antibiotic treated animals. It also suggests that residual mycobacterial antigens may be important for protection. This study establishes a viable experimental model for investigating the immune factors (such as depletion of cell types or neutralization of cytokines) that play a role in concomitant immunity against Mtb in NHP.

## Supporting information

NHPdata

MtbBarcodingData

## Acknowledgements

This study was funded by NIH R01 AI114674 (Flynn/Fortune), NIH NIAID grant 75N93019C00071 IMPAcTB (Flynn/Fortune) and the Bill and Melinda Gates Foundation. We are grateful to our veterinary and research technicians for the dedication and work on this study. We acknowledge H. Jacob Borish and Alexander White for assistance with PET CT analyses and Dr. Charles Scanga for project management.

## Methods

### Ethics Statement

All experimental manipulations, protocols, and care of the animals were approved by the University of Pittsburgh School of Medicine Institutional Animal Care and Use Committee (IACUC). The protocol assurance number for our IACUC is A3187-01. Our specific protocol approval numbers for this project are 15066174 and 18124275. The IACUC adheres to national guidelines established in the Animal Welfare Act (7 U.S.C. Sections 2131–2159) and the Guide for the Care and Use of Laboratory Animals (8th Edition) as mandated by the U.S. Public Health Service Policy.

All macaques used in this study were housed at the University of Pittsburgh in rooms with autonomously controlled temperature, humidity, and lighting. Animals were singly housed in caging at least 2 square meters apart that allowed visual and tactile contact with neighboring conspecifics. The macaques were fed twice daily with biscuits formulated for nonhuman primates, supplemented at least 4 days/week with large pieces of fresh fruits or vegetables. Animals had access to water *ad libitem*. Because our macaques were singly housed due to the infectious nature of these studies, an enhanced enrichment plan was designed and overseen by our nonhuman primate enrichment specialist. This plan has 3 components. First, species-specific behaviors are encouraged. All animals have access to toys and other manipulata, some of which will be filled with food treats (e.g. frozen fruit, peanut butter, etc.). These are rotated on a regular basis. Puzzle feeders, foraging boards, and cardboard tubes containing small food items also are placed in the cage to stimulate foraging behaviors. Adjustable mirrors accessible to the animals stimulate interaction between animals. Second, routine interaction between humans and macaques are encouraged. These interactions occur daily and consist mainly of small food objects offered as enrichment and adhere to established safety protocols. Animal caretakers are encouraged to interact with the animals (by talking or with facial expressions) while performing tasks in the housing area. Routine procedures (e.g. feeding, cage cleaning, etc) are done on a strict schedule to allow the animals to acclimate to a routine daily schedule. Third, all macaques are provided with a variety of visual and auditory stimulation. Housing areas contain either radios or TV/video equipment that play cartoons or other formats designed for children for at least 3 hours each day. The videos and radios are rotated between animal rooms so that the same enrichment is not played repetitively for the same group of animals.

All animals are checked at least twice daily to assess appetite, attitude, activity level, hydration status, etc. Following *M. tuberculosis* infection, the animals are monitored closely for evidence of disease (e.g., anorexia, weight loss, tachypnea, dyspnea, coughing). Physical exams, including weights, are performed on a regular basis. Animals are sedated prior to all veterinary procedures (e.g. blood draws, etc.) using ketamine or other approved drugs. Regular PET CT imaging is conducted on most of our macaques following infection and has proved very useful for monitoring disease progression. Our veterinary technicians monitor animals especially closely for any signs of pain or distress. If any are noted, appropriate supportive care (e.g. dietary supplementation, rehydration) and clinical treatments (analgesics) are given. Any animal considered to have advanced disease or intractable pain or distress from any cause is sedated with ketamine and then humanely euthanatized using sodium pentobarbital.

### Animals, infections, drug treatment and disease tracking by PET CT

Twenty cynomolgus macaques (*Macaca fascicularis*) with age range of 5.3-9.1 years were obtained from Valley Biosystems (3 females, 17 males, Sacramento, California). All animals were placed in quarantine for 1 month where they were monitored to ensure good physical health and no prior Mtb infection. Fourteen animals were infected with DNA-tagged Mtb Erdman (Library P) via bronchoscopic instillation (4-16 CFU) as previously described^23^. Six macaques did not receive Library P (naïve controls). Granuloma formation, lung inflammation and overall disease in the Library P infected macaques was tracked using ^18^F-fluorodeoxyglucose (FDG) PET CT every 4 weeks. PET CT scans were analyzed using OsiriX viewer as previously described with a detection limit of 1mm^24^. After 16 weeks of infection, 7 animals were treated with antibiotics and the other 7 animals were not (no-Abx). Because this study was scheduled to last for 9 months, we needed to choose macaques from the first cohort of 10 animals for the non-drug treated group that we predicted would survive until the end of the study without drug intervention. The 3 macaques were chosen based on minimal change in granuloma numbers from 4 to 12 weeks post-infection and reduced total lung inflammation measured by PET CT. We previously described that lack of new granuloma formation and a stable metabolic activity measured by FDG in granulomas and lymph nodes early post-infection could predict which macaques will develop controlled or “latent” infection at 6 months post-infection^24^. Macaques in the drug-treated group were given antibiotics orally once daily for ∼3 months (80-97 days) (Rifampicin 20mg/kg; Isoniazid 15mg/kg; Ethambutol 50mg/kg; Pyrazinamide 150mg/kg)^25^. Compliance ranged from 80-100% although one macaque had low compliance (Animal number 30416, 62%). Macaques were taken off drug treatment 2-4 weeks before the second infection with Mtb library S and subsequently necropsied 4 weeks later. The macaques in the non-drug treated group followed the same timeline as the drug-treated group but without drug treatment. All twenty macaques (14 Library P infected and six naïve macaques) then were challenged with 7-22 CFU of Mtb library S and necropsied 4 weeks later. Dose was calculated from colony counts after plating an aliquot of the infection inoculum on 7H11 agar plates and incubating for 3 weeks at 37°C/5% CO_2_.

### Bronchoalveolar lavage (BAL)

BAL was obtained every 8 weeks as described previously^26,27^. Briefly, a bronchoscope with a 2.5 mm outer diameter was inserted into the trachea of a sedated animal and placed in the right middle or lower lobe. A saline solution (40ml) was introduced briefly, suctioned, and transferred to a 50 ml conical tube. An aliquot was used to plate for CFU on 7H11 agar which was read after 3 weeks of incubation at 37°C/5% CO_2_. BAL were centrifuged at 1,800 rpm for 8 minutes at 4°C. Cells were resuspended in 1ml PBS, counted using a hemocytometer and used for intracellular cytokine staining. Supernatants were filtered with 0.22 μm syringe filter to remove Mtb bacteria and frozen at −80°C until use for Luminex.

### Necropsy Procedures

Procedures done during necropsy have been previously described^23,27^. Briefly, 1-3 days prior to necropsy, a PET CT scan was taken and used to identify the location and metabolic activity (FDG activity) of granulomas and lymph nodes; this scan was used as a map to aid in the individual identification and excision of these samples during necropsy. On the day of necropsy, macaques were humanely sacrificed with sodium pentobarbital injection and terminally bled. Individual granulomas, thoracic and peripheral lymph nodes, lung tissue, spleen and liver were all excised and homogenized separately into single cell suspensions. Homogenates were aliquoted for plating on 7H11 agar for bacterial burden, freezing for DNA extraction and staining for flow cytometry analysis. Any remaining samples were frozen for future use. Homogenates were plated in serial dilutions on 7H11 medium and incubated at 37°C/5% CO_2_ for 3 weeks before enumeration of CFU.

### Isolation of genomic DNA from bacteria

DNA extraction was performed on granuloma and lymph node homogenates, as well as their scrapates (scraped colonies that grew on 7H11 agar plates) for library identification as described previously^14^. Briefly, a small aliquot of the homogenate or scrapate were vortexed with 0.1mm zirconia-silica beads (BioSpec Products, Inc.) and subsequently extracted twice with phenol chloroform isoamyl alcohol (25:24:1, Sigma-Aldrich) before precipitating DNA with molecular grade 100% isopropanol (Sigma-Aldrich) and 3M sodium acetate (Sigma-Aldrich) and resuspending in nuclease-free water (Invitrogen).

### Library identification

Identification of library DNA tags have been previously described^13^. Briefly, DNA was amplified by PCR for 24-36 cycles before using in the NanoString nCounter assay (NanoString Technologies) with custom designed probes. The scheme for labeling granulomas as old or new is found in **Table 1**.

**Table 1.** Granuloma Classification scheme.

| PET CT* | Library | Classification |
| --- | --- | --- |
| old | P | old |
| old | S | new |
| new | P | old |
| new | S | new |
| old/new | P + S | new |
| unknown | P | old |
| unknown | S | new |
| old | No data | old |
| new | No data | new |

\* granuloma was noted on PET CT scan during first infection (old) or only after second infection (new). Unknown: granuloma was not seen on PET CT scan, possibly due to size or inability to discriminate from other granulomas in close proximity.

### Intracellular cytokine staining and flow cytometry

Intracellular cytokine staining was performed on BAL and individual granuloma samples. BAL cells (250,000-1x10^6^) were stimulated with peptide pools of ESAT-6 and CFP-10 (10μg/ml of each peptide pool) in the presence of Brefeldin A (GolgiPlug, BD Biosciences) at 37°C/5% CO_2_ for 3.5-4 hours prior to staining. Unstimulated controls were always included, however, positive controls (phorbol dibutyrate [PdBu] and ionomycin were only included if there were enough cells^27^. Cells were stained with a viability marker (LIVE/DEAD fixable blue dead cell stain kit, Invitrogen) and surface and intracellular markers. Surface markers for T cells include CD3 (clone SP34-2, BD Pharmingen), CD4 (Clone L200, BD Horizon) and CD8 (Clone SK1, BD Biosciences). Intracellular markers include IFNγ (Clone B27, BD Biosciences), TNF (Clone MAB11, BD Biosciences), IL-2 (Clone MQ1-17H12, Biolegend), IL-10 (Clone JES3-9D7, eBioscience) and Ki67 (Clone B56, BD Biosciences). Data was acquired using the LSR II (BD) and analyzed using FlowJo software v10.6.1 (BD).

Because of the abundance of Mtb antigens already present in granulomas and involved lymph nodes^28^, these samples were not further stimulated with Mtb peptides. All samples were incubated in the presence of Brefeldin A (GolgiPlug, BD Biosciences) at 37°C/5% CO_2_ for 3.5-4 hours prior to staining. Cells were stained with a viability marker (LIVE/DEAD fixable blue dead cell stain kit, Invitrogen) and surface and intracellular markers. Surface markers include CD3 (clone SP34-2, BD Pharmingen), CD4 (Clone L200, BD Horizon), CD8 (Clone SK1, BD Biosciences) and CD20 (Clone 2H7, eBioscience). Intracellular markers include CD107a (Clone H4A3, Biolegend), IFNγ (Clone B27, BD Biosciences), TNF (Clone MAB11, BD Biosciences), IL-2 (Clone MQ1-17H12, Biolegend), IL-10 (Clone JES3-9D7, eBioscience) and IL-17 (Clone eBio64CAP17, eBioscience). Data was acquired using the LSR II (BD) and analyzed using FlowJo software v10.6.1 (BD).

### Luminex on BAL samples

Frozen supernatants were thawed on ice and concentrated using a centrifugation filter unit (Amicon Ultra-4 Centrifugal Filter Unit, 3kDa cutoff, Millipore Sigma) following manufacturer’s instructions. Samples were assayed in duplicates using the ProcartaPlex NHP multiplex immunoassay (Invitrogen) following manufacturer’s instructions measuring levels of 30 cytokines and chemokines using the BioPlex reader (Biorad). An additional 2-fold dilution was performed on the supplied standards to extend its lower detection limit.

### Statistical Analysis

We used D’Agostino & Pearson test to determine normality of data, but due to small sample sizes, nonparametric tests were used. The Mann-Whitney test was used to compare two groups. The Kruskal-Wallis test with Dunn’s adjustment for multiple comparisons was used to compare three or more groups. P-values < 0.05 were considered statistically significant. Data were graphed and analyzed using GraphPad Prism v10 (GraphPad Software). All CFU data in graphs with the y-axis in log scale were transformed by adding 1.

## Data availability statement

The data used to generated the accompanying figures is provided in Supplementary Tables 1 and 2.

## References

1. Cox, F. E. Concomitant infections, parasites and immune responses. Parasitology 122 Suppl, S23-38 (2001).

2. Andrews, J. R. et al. Risk of progression to active tuberculosis following reinfection with Mycobacterium tuberculosis. Clin Infect Dis 54, 784–791 (2012).

3. Belkaid, Y., Piccirillo, C. A., Mendez, S., Shevach, E. M. & Sacks, D. L. CD4+CD25+ regulatory T cells control Leishmania major persistence and immunity. Nature 420, 502–507 (2002).

4. Behr, M. A., Kaufmann, E., Duffin, J., Edelstein, P. H. & Ramakrishnan, L. Latent Tuberculosis: Two Centuries of Confusion. Am J Respir Crit Care Med 204, 142–148.

5. He, W. et al. Endogenous relapse and exogenous reinfection in recurrent pulmonary tuberculosis: A retrospective study revealed by whole genome sequencing. Front Microbiol 14, 1115295 (2023).

6. Micheni, L. N., Kassaza, K., Kinyi, H., Ntulume, I. & Bazira, J. Detection of Mycobacterium tuberculosis multiple strains in sputum samples from patients with pulmonary tuberculosis in south western Uganda using MIRU-VNTR. Sci Rep 12, 1656 (2022).

7. Micheni, L. N., Deyno, S. & Bazira, J. Mycobacterium tuberculosis mixed infections and drug resistance in sub-Saharan Africa: a systematic review. Afr Health Sci 22, 560–572 (2022).

8. Henao-Tamayo, M. et al. A mouse model of tuberculosis reinfection. Tuberculosis (Edinb) 92, 211–217 (2012).

9. Nemeth, J. et al. Contained Mycobacterium tuberculosis infection induces concomitant and heterologous protection. PLoS Pathog 16, e1008655 (2020).

10. Shen, X. et al. Recurrent tuberculosis in an urban area in China: relapse or exogenous reinfection? Tuberculosis (Edinb) 103, 97–104 (2017).

11. Millet, J.-P. et al. Tuberculosis recurrence and its associated risk factors among successfully treated patients. J Epidemiol Community Health 63, 799–804 (2009).

12. Verver, S. et al. Rate of reinfection tuberculosis after successful treatment is higher than rate of new tuberculosis. Am J Respir Crit Care Med 171, 1430–1435 (2005).

13. Cadena, A. M. et al. Concurrent infection with Mycobacterium tuberculosis confers robust protection against secondary infection in macaques. PLoS Pathog 14, e1007305 (2018).

14. Lin, P. L. et al. Sterilization of granulomas is common in active and latent tuberculosis despite within-host variability in bacterial killing. Nat Med 20, 75–79 (2014).

15. Joslyn, L. R., Flynn, J. L., Kirschner, D. E. & Linderman, J. J. Concomitant immunity to M. tuberculosis infection. Sci Rep 12, 20731 (2022).

16. Loetscher, M. et al. Chemokine receptor specific for IP10 and mig: structure, function, and expression in activated T-lymphocytes. Journal of Experimental Medicine 184, 963–969 (1996).

17. Vissers, J. L., Hartgers, F. C., Lindhout, E., Figdor, C. G. & Adema, G. J. BLC (CXCL13) is expressed by different dendritic cell subsets in vitro and in vivo. Eur J Immunol 31, 1544– 1549 (2001).

18. Galeano Niño, J. L., et al. Cytotoxic T cells swarm by homotypic chemokine signalling. eLife 9, e56554 (2020).

19. Darrah, P. A. et al. Prevention of tuberculosis in macaques after intravenous BCG immunization. Nature 577, 95–102 (2020).

20. Peters, J. M. et al. Protective intravenous BCG vaccination induces enhanced immune signaling in the airways. 2023.07.16.549208 Preprint at 10.1101/2023.07.16.549208 (2023).

21. Luzze, H. et al. Relapse more common than reinfection in recurrent tuberculosis 1-2 years post treatment in urban Uganda. Int J Tuberc Lung Dis 17, 361–367 (2013).

22. Marx, F. M. et al. The temporal dynamics of relapse and reinfection tuberculosis after successful treatment: a retrospective cohort study. Clin Infect Dis 58, 1676–1683 (2014).

23. Lin, P. L. et al. Quantitative comparison of active and latent tuberculosis in the cynomolgus macaque model. Infect Immun 77, 4631–4642 (2009).

24. White, A. G. et al. Analysis of 18FDG PET/CT Imaging as a Tool for Studying Mycobacterium tuberculosis Infection and Treatment in Non-human Primates. J Vis Exp 56375 (2017) doi:10.3791/56375.

25. Lin, P. L. et al. Metronidazole prevents reactivation of latent Mycobacterium tuberculosis infection in macaques. Proc Natl Acad Sci U S A 109, 14188–14193 (2012).

26. Capuano, S. V. et al. Experimental Mycobacterium tuberculosis infection of cynomolgus macaques closely resembles the various manifestations of human M. tuberculosis infection. Infect Immun 71, 5831–5844 (2003).

27. Lin, P. L. et al. Early events in Mycobacterium tuberculosis infection in cynomolgus macaques. Infect Immun 74, 3790–3803 (2006).

28. Gideon, H. P. et al. Variability in tuberculosis granuloma T cell responses exists, but a balance of pro-and anti-inflammatory cytokines is associated with sterilization. PLoS Pathog 11, e1004603 (2015).

