## SupplementaryFigures for "Antibiotic treatment modestly reduces protection against Mycobacterium tuberculosis reinfection in macaques"

**Supplementary Figures:**

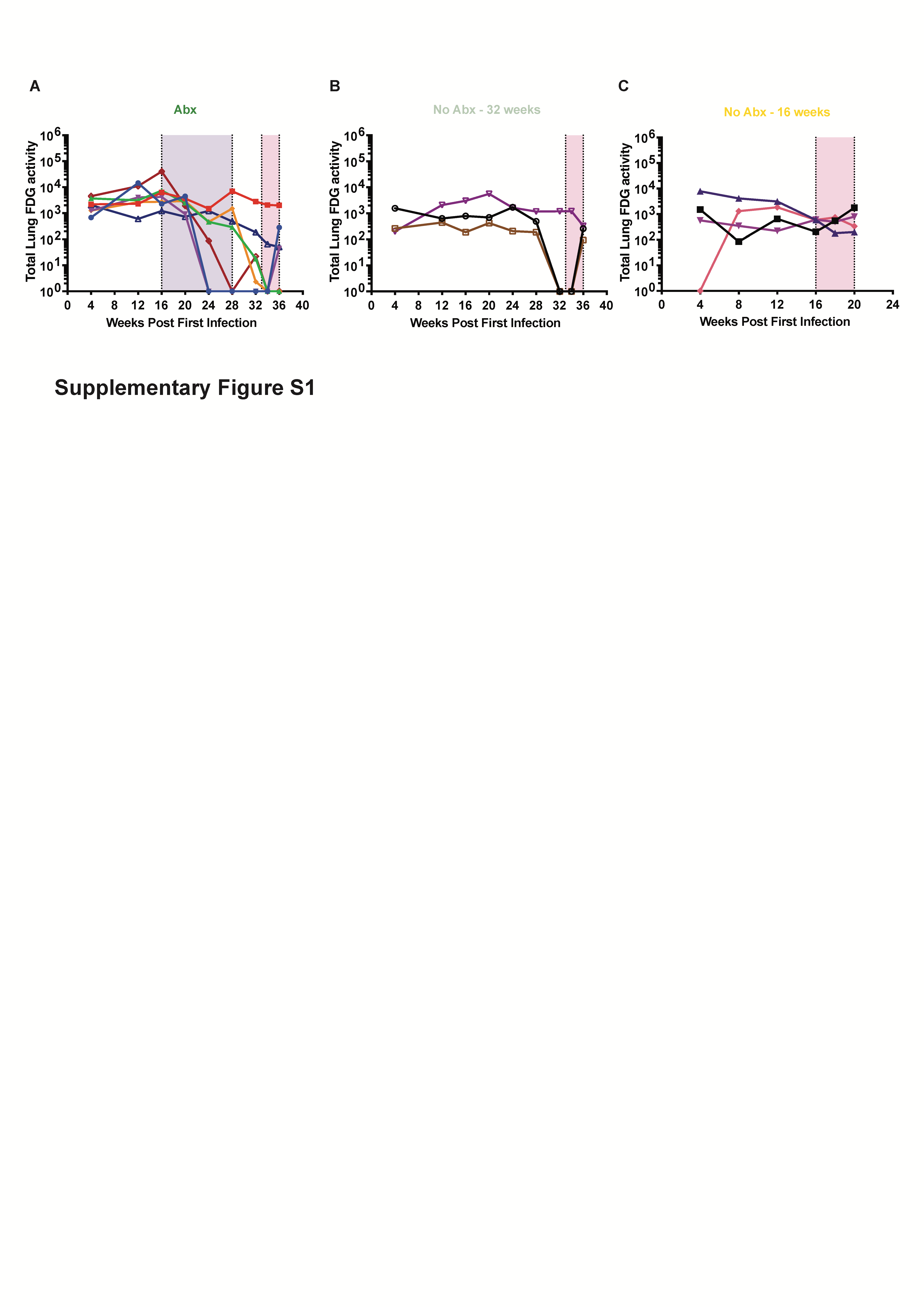

**Figure S1. Total lung PET-CT FDG activity reflects inflammatory changes over the course of the experiment.** Serial FDG PET CT scans were performed throughout the primary and secondary challenge. Total lung FDG activity is shown for the Abx (A) and the two no-Abx (B, C) groups. Purple shaded area indicates duration of drug treatment and pink shaded area reflects the time and duration of the secondary challenge with Mtb.
